# Identification of slow-cycling germline stem cells and their regulation by PLZF

**DOI:** 10.1101/477604

**Authors:** Manju Sharma, Anuj Srivastava, Heather E. Fairfield, David Bergstrom, William F. Flynn, Robert E Braun

**Affiliations:** The Jackson Laboratory, 600 Main Street, Bar Harbor, ME, 04609, USA

**Keywords:** Spermatogonial stem cell, SSC, EOMES, GDNF, GFRA1, PLZF

## Abstract

Long-term maintenance of spermatogenesis in mammals is supported by GDNF, an essential growth factor required for spermatogonial stem cell (SSC) self-renewal. Exploiting a transgenic GDNF overexpression model, which expands and normalizes the pool of undifferentiated spermatogonia between *Plzf*^+/+^ and *Plzf*^*lu/lu*^ mice, we used RNAseq to identify a rare subpopulation of cells that express EOMES, a T-box transcription factor. Lineage tracing and busulfan challenge show that these are long-term SSCs that contribute to steady state spermatogenesis as well as regeneration following chemical injury. EOMES+ SSCs have a lower proliferation index than EOMES− GFRA1+ spermatogonia in wild-type but not in *Plzf*^*lu/lu*^ mice. This comparison demonstrates that PLZF regulates their proliferative activity and suggests that EOMES+ SSCs are lost through proliferative exhaustion in *Plzf*^*lu/lu*^ mice. Single cell RNA sequencing of EOMES+ cells from *Plzf*^+/+^ and *Plzf*^*lu/lu*^ mice support a hierarchical model of both slow- and rapid-cycling SSCs.

## INTRODUCTION

Fertility in males is supported by a robust stem cell system that allows for continuous sperm production throughout the reproductive life of the individual. In humans this lasts for decades and in a mouse can last for nearly its entire lifetime. However, despite more than a half century of research, and intensive investigation by many labs over the last decade, the identity of the germline stem cell continues to be elusive and controversial.

Stem cell function clearly resides in a subpopulation of spermatogonia within the basal compartment of the seminiferous tubules. In 1971, Huckins and Oakberg proposed the “A_s_” model of spermatogonial stem cells (SSC) function where A_single_ (A_s_) SSCs divide in a linear and non-reversible manner to populate the spermatogenic lineage (*1–3*). In this model, A_s_ spermatogonia are the SSCs, dividing symmetrically and with complete cytokinesis to form two daughter A_s_ cells for self-renewal, or dividing with incomplete cytokinesis to form A_paired_ (A_pr_) cells, which are irreversibly committed to differentiation. A_pr_ cells, in turn, divide to form A_aligned_ (A_al_) spermatogonia, which exist as chains of 4, 8, or 16 interconnected cells. The A_s_, A_pr_, and A_al_ spermatogonia encompass the pool of undifferentiated spermatogonia that can be identified morphologically and are thought to share important functional properties distinct from the differentiated spermatogonia; A_1_-A_4,_ intermediate and B (*4, 5*).

Contrary to the Huckins/Oakberg A_s_ model, recent studies on the behavior of pulse-labeled spermatogonia populations suggest that A_pr_ and A_al_ syncytia can fragment and revert to become A_s_ cells following transplantation as well as during steady state spermatogenesis (*6, 7*). This suggests that undifferentiated spermatogonia are not irreversibly committed to differentiation, allowing for an alternative mechanism for SSC self-renewal. Furthermore, A_s_, A_pr_, and A_al_ spermatogonia are characterized by heterogeneous gene expression (*6–11*) including recent descriptions of specific subpopulations of A_s_ cells with SSC activity expressing *Id4*, *Pax7*, *Bmi1* and *T* (*12–16*) (*17*). These new data do not easily comport to a unifying model and imply that the mode of SSC function in the testes is more complex than the original Huckins-Oakberg A_s_ model suggests.

A_s_ and A_pr_ cells express GFRA1, a GPI-anchored receptor for glial cell-derived neurotrophic factor (GDNF) (*18–22*). GDNF is secreted by neighboring somatic Sertoli (*23*) and peritubular myoid (*24*) cells and is required for establishment and self-renewal of the SSC population in a dose-dependent manner (*23*). A decrease in GDNF levels results in germ cell loss, while overexpression of GDNF promotes accumulation of SSCs due to a block in differentiation (*23, 25*). PLZF, expressed in A_s_, A_pr_ and A_al_ spermatogonia, is a transcription factor required for SSC maintenance, as mutation of *Plzf* results in age-dependent germ cell loss (*26, 27*). The mechanisms by which PLZF regulates SSC maintenance are not yet known. We describe here the identification of a rare subpopulation of A_s_ cells, which are slow-cycling and regulated by PLZF.

## RESULTS

### GDNF increases the undifferentiated spermatogonial population in *Plzf* mutants

Stage-specific temporal ectopic expression of GDNF in supporting Sertoli cells results in the accumulation of large clusters of tightly-packed PLZF+ undifferentiated spermatogonia (*25*). To determine whether overexpression of GDNF could rescue germ cell loss in the *luxoid* (*lu*) mutant, we generated *Tg(Ctsl-Gdnf)*^*1Reb*^; *Plzf*^*lu/lu*^ mice (referred to as *Tg(Gdnf);lu/lu*). While *lu/lu* mice are smaller than wild-type (WT), we found no significant difference in body weight between age-matched *lu/lu* and *Tg(Gdnf);lu/lu* mice (Fig. 1A). However, at four months of age, testis weight was significantly higher in *Tg(Gdnf);lu/lu* mice compared to *lu/lu* (p<0.001), although it was still lower than in *Tg(Gdnf)* (p<0.0001) animals (Fig. 1B). Periodic acid-Schiff staining of testes sections showed fewer agametic tubules (referred to as Sertoli cell only) in *Tg(Gdnf);lu/lu* mice compared to *lu/lu* at both 4 and 6 months of age (Fig. 1C and D). Increased testis weight in *Tg(GDNF);lu/lu* mice could therefore be due to an increase in the number of cells occupying individual tubules, reflected by a decrease in the number of tubules with a Sertoli cell only phenotype, and fewer tubules with loss of one or more cell populations at 6 months of age (Fig 1D).

**Figure 1.**
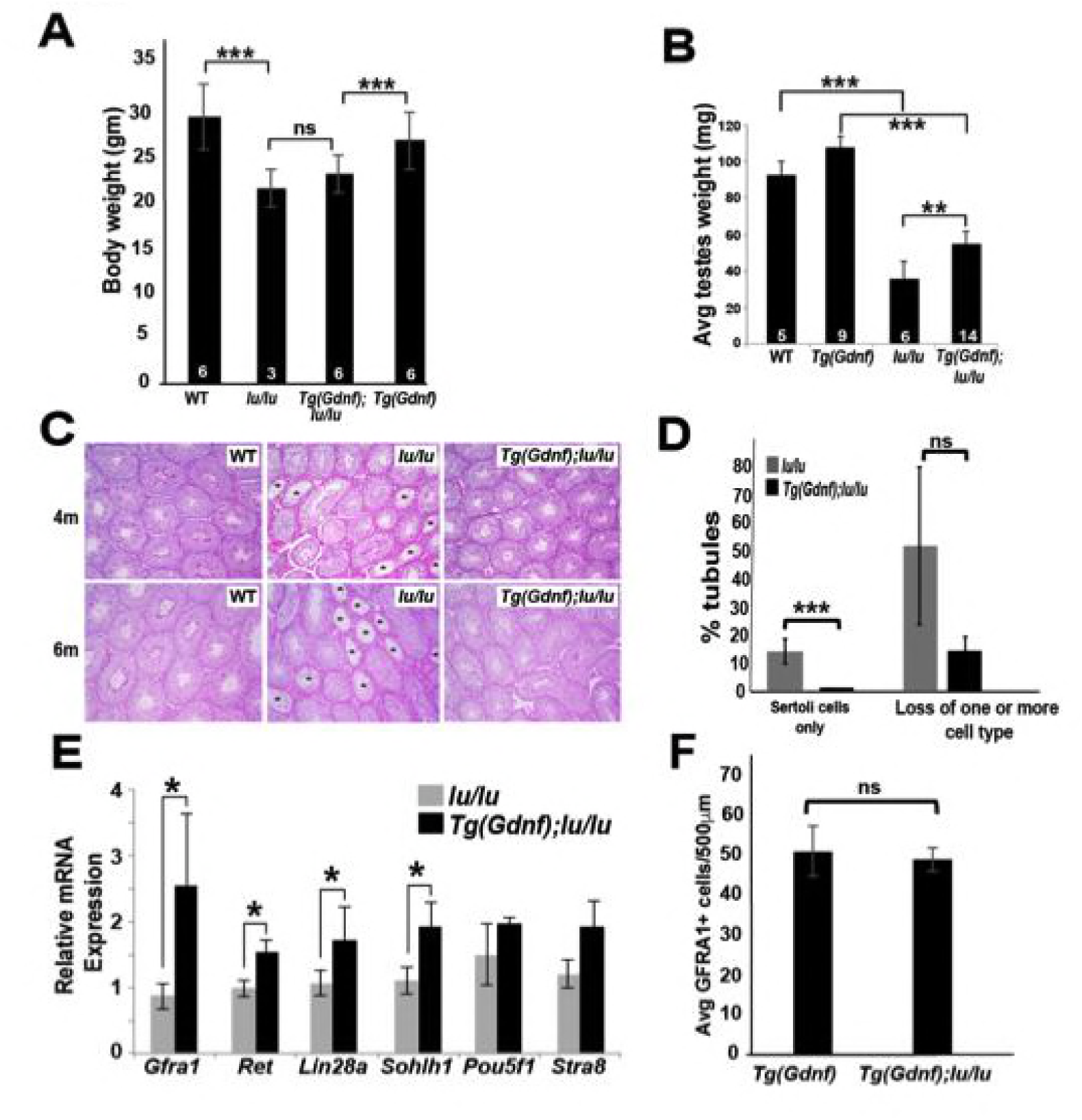
Sertoli-cell overexpression of GDNF partially reverses the loss of undifferentiated spermatogonia in *luxoid* mice. (A) No difference was observed in body weight between *lu*/*lu* and *Tg*(*Gdnf)*;*lu*/*lu* male mice at 4 months of age. ns, not significant; ***, p <0.005; number in bar = n animals. (B) Significant differences were observed in testis weight between *lu*/*lu* and *Tg(Gdnf);lu/lu* animals. **, p<0.005; ***, p<0.0001; number in bar = n testes. (C) Representative images of periodic-acid-Schiff stained cross-sections of 4- and 6-month old testes showing agametic tubules (asterisks) in *lu*/*lu*. (D) Fewer tubules with germ cell loss were present at 6 months in *Tg(Gdnf):lu/lu*. A total of 200-300 tubules were counted for each genotype (n=3). P < 0.005. (E) qRT-PCR on testis RNA from 4-month old mice shows a significant increase in SSC gene expression in *lu/lu* mice overexpressing GDNF in Sertoli cells. Relative mRNA levels are normalized to a *β-actin* internal control. *, p<0.01. (n = 3) (F) Average number of GFRA1+ cells per 500μm length of tubules in *Tg(Gdnf)* and *Tg(Gdnf);lu/lu* in 10wk old mice (n=3).

To determine what germ cell populations were expanded in *Tg(Gdnf);lu/lu* testes, we immuno-stained both whole-mount tubules and sections for spermatogonia markers. Large clusters of GFRA1+ cells were observed in *Tg(Gdnf);lu/lu* tubules, suggesting an increased number of the earliest spermatogonia cell types, A_s_ and A_pr_ (Fig. S1A). Quantitation of LIN28A staining, which is detected in a subpopulation of A_s_, and A_pr_, A_al_ and A1 spermatogonia (*28*), on 12-week old testes sections (Fig. S1B) showed that 75% of *Tg(Gdnf);lu/lu* tubule cross-sections had >5 LIN28A+ cells per tubule compared to 30% in *lu/lu* (Fig. S1C, p<0.001). Only 8% of tubules lacked LIN28A-labeled cells compared to 30% in *lu/lu* testes (p<0.01) (Fig. S1C). To determine whether the A_1_ to A_4_ differentiated spermatogonia population increased in *Tg(Gdnf);lu/lu*tubules, we immunostained for the marker SOHLH1, which is not expressed in PLZF+ A_s_ to A_al_ cells (Fig. S1D). Similar to LIN28A, quantification of sectioned tubules revealed a significant increase in the percent of tubules with >5 SOHLH1+ cells (Fig. S1E, F).

Using quantitative reverse transcriptase PCR (qRT-PCR), we compared mRNA levels of several known genes expressed in undifferentiated spermatogonia, using total RNA from 4 month-old *lu/lu* and *Tg(Gdnf);lu/lu* testes. Similar to our immunostaining results, we observed a significant increase in the undifferentiated spermatogonia markers *Gfra1*, *Lin28a* and *Sohlh1* in *Tg(Gdnf);lu/lu* compared to *lu/lu* (Fig. 1E). We also saw an increase in *Ret*, a gene that is part of the GDNF-GFRA1 complex. Thus, temporal ectopic expression of GDNF significantly increases the numbers of tubules with both undifferentiated and differentiated spermatogonia, leading to a partial rescue of testis weight in *lu/lu* mice.

### Unbiased discovery of genes whose expression is altered in *Plzf* mutants

Seminiferous tubules from *Tg(Gdnf)* mice have an expanded GFRA1+ population and enhanced ability to repopulate the spermatogonial lineage in testes transplants compared to WT (*25*). Because *Tg(Gdnf)* and *Tg(Gdnf);lu/lu* testes have a comparable number of GFRA1+ spermatogonia (Fig. 1F), we reasoned that these mice could be used as a valuable tool for the discovery of genes involved in self-renewal. Using RNAseq, we compared the transcriptomes of whole testes from *Tg(Gdnf)* and *Tg(Gdnf);lu/lu* mice and searched for genes with significant fold-changes in expression. Of 20,000 gene transcripts, ~1,500 were differentially expressed with an absolute log_2_ fold change >1.5 (FDR<0.05). Of this subset, 150 were upregulated two-fold or more, and 60 were down-regulated two- to seven-fold in *luxoid* mutants.

As expected, many previously known gene transcripts specific to undifferentiated spermatogonia had no significant change in expression between genotypes including: *Gfra1*, *Ret, Lin28a, Utf1, Nanos2, Nanos3, Sohlh2,* and *Sall4* (Fig. 2A), confirming that the undifferentiated spermatogonial populations were similar between the two genotypes. In the group of genes that were significantly down-regulated in *Tg(Gdnf);lu/lu* testes, we identified a number of genes that are involved in the maintenance of pluripotent stem cell fate or early embryonic development (Fig. 2B) including: *Pou5f1*, *Brachyury (T)* and *Eomes*. *Pou5f1*, essential for maintaining pluripotency of ES cells and for maintaining viability of the mammalian germline (*29, 30*), was down-regulated two-fold in *luxoid* testes overexpressing GDNF. *Brachyury (T)*, down-regulated seven-fold in *Tg(Gdnf);lu/lu* testes, is a T-box transcription factor important for germ cell development that when overexpressed results in testicular germ-cell tumors (*31, 32*). *Eomes*, a T-box transcription factor that was down-regulated 3-fold in *Tg(Gdnf);lu/lu* testes, is known to play a critical role in embryonic development (*33, 34*). Notably, the read counts per million (RCPM) for the two highly down-regulated genes shown, *Eomes* and *T*, were very low in either genotype and found at the distant end of the expression spectrum amongst 20,000 transcripts (Fig. 2C, far right). *Eomes* had a value of only 1.4 RCPM in *Tg(Gdnf)* testis RNA compared to 5,509 RCPM for *Prm2*, a gene expressed in spermatids. This suggests that *Eomes* and *T* are targets of PLZF and are down-regulated in *lu/lu* mice, or that T and EOMES are expressed in a small subset of germ cells that are specifically reduced in the *lu/lu* mutant, possibly within a restricted population of A_s_ cells.

**Figure 2.**
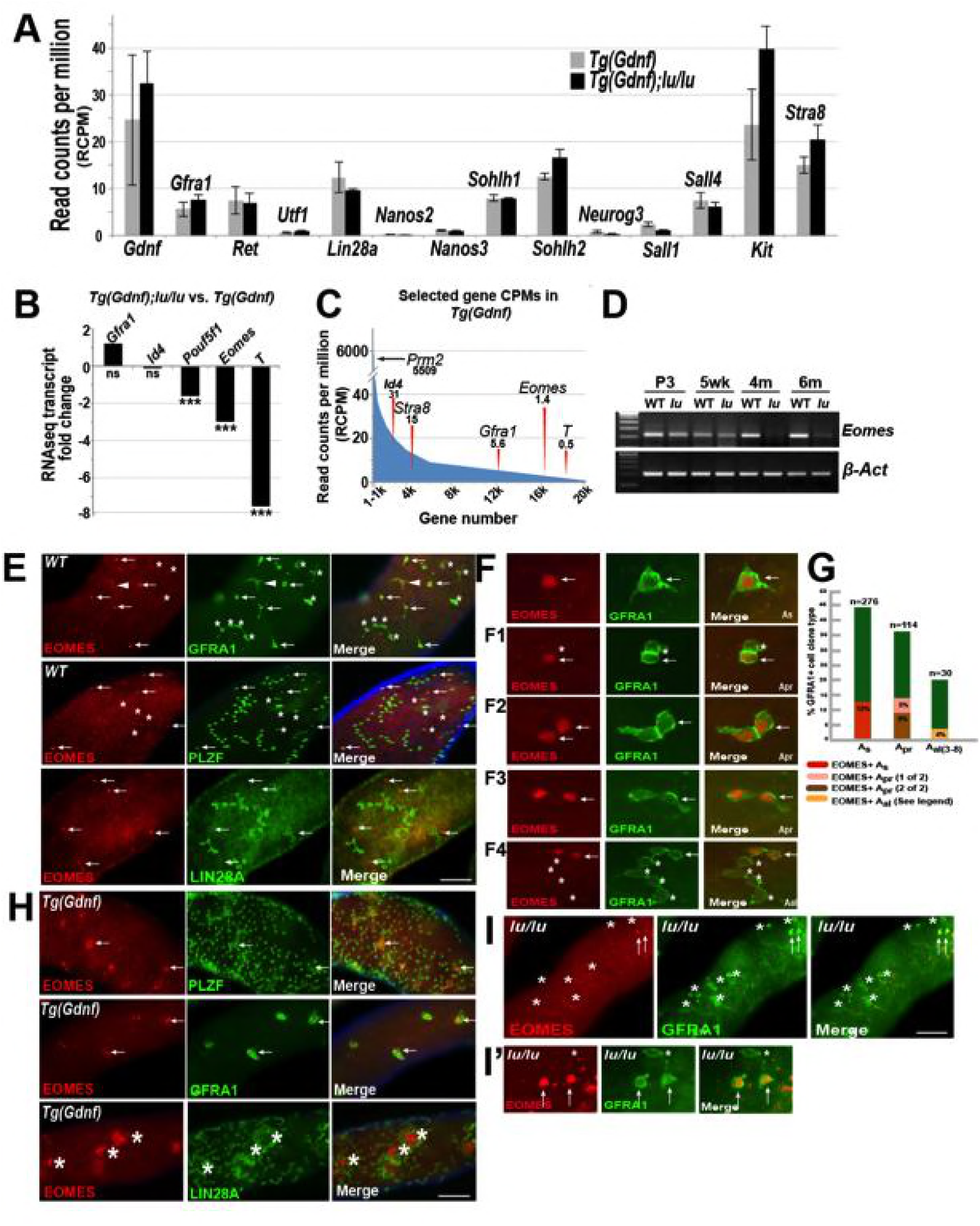
EOMES marks a subpopulation of SSCs. (A) Relative gene expression (read counts per million, RCPM) of selected transcripts from RNAseq performed on testis RNA from 4-month old *Tg(Gdnf)* and *Tg(Gdnf);lu/lu* mice. No significant change is detected in known spermatogonial marker genes. (B) Fold change in RNAseq transcripts for a subset of pluripotency and developmental genes in *Tg(Gdnf);lu/lu* compared to *Tg*(*Gdnf)* testes. ***, p<0.0001. (C) Relatively low RCPM of the highest-fold change transcripts shown in (B), compared to the spermatid-expressed protamine 2 gene *Prm2*, and differentiating spermatogonia marker *Stra8*. X-axis represents 20,000 genes numbered in descending order by RCPM. (D) Semi-quantitative RT-PCR of total testis RNA showing decreasing *Eomes* expression over time in *lu*/*lu* testes. (E) Immunostaining for EOMES and undifferentiated spermatogonial markers in WT whole-mount testes. EOMES marks some (arrows), but not all (asterisks), GFRA1+ PLZF+ A_s_ cells. Weak EOMES staining can occasionally be detected in chains of GFRA1+ A_al_ cells (arrowhead). EOMES+ A_s_ cells do not express LIN28A (bottom row, arrows). (F) Examples of EOMES+ subset of GFRA1+ cells. F1 shows an example of two adjacent cells where one is EOMES+ (arrow) and one is EOMES-(asterisk). F2 and F2 illustrate two examples of two juxtaposed cells that are both EOMES+. F4 illustrates an example of a chain of 6 A_al_ cells with two EOMES+ (arrow) and four EOMES– (asterisks) cells. (G) Graph showing the percent of EOMES+ cell in the GFRA1+ population clone types in WT adult mice (631 GFRA1+ cells from 3 mice). n = number of clone types counted (e.g. A_pr_ = 114 pairs of cells = 228 GFRA1+ total cells). Of the total GFRA1+ A_pr_ population, in 9% of cases both cells were positive for EOMES, while 5% had only one cell positive for EOMES. Within the A_al_ fraction, 4% of the clones express EOMES in a heterogeneous pattern (two A_al_ chains where 3 of 3 were EOMES+; two A_al_ chains of 4 where all were EOMES+; one A_al_ chain of 4 where only 1 was EOMES+; one A_al_ chain of 6 where only 2 of 6 cells were EOMES+). (H) EOMES is expressed in tightly packed GFRA1+ PLZF+ clusters (arrows) found in *Tg*(*Gdnf)* whole-mount tubules. EOMES is not expressed in the LIN28A+ cells in the cortical region of the clusters (bottom row, asterisks). (I) Detection of EOMES+ GFRA1+ cells (arrows) in *Plzf ^lu/u^* mutants. Asterisks mark an EOMES-GFRA1+ cell.

We next validated candidate genes at different developmental time points by RT-PCR. Using total testis RNA from *Tg(Gdnf)* and *Tg(Gdnf);lu/lu* mice, *Eomes* was reduced in *lu/lu* testes as early as 3 days after birth, and became barely detectable at 4 and 6 months of age (Fig. 2D). As mutations in *Plzf* lead to age-dependent germ cell loss, *Eomes* was a compelling candidate marking a functional SSC pool that is depleted in *luxoid* mutants.

### EOMES is detected in a subpopulation of GFRA1+ cells in the testis

We first determined whether EOMES was expressed within the spermatogonia population of the testis. We immuno-stained whole mount WT seminiferous tubules for EOMES together with GFRA1, which marks A_s_, A_pr_ and small chains of A_al_ spermatogonia (Fig. 2E, F, arrows). EOMES was expressed in 12% of the total GFRA1+ A_s_ population of cells (Fig. 2G). Fourteen percent of GFRA1+ A_pr_ clones also expressed EOMES+ in either one (5%) or both (9%) of the cells (Fig. 2F1, 2F2, and 2F3, arrows, and G). The clones counted as A_pr_ could be two lineage-independent, but adjacent, A_s_ cells, A_pr_ cells about to complete cytokinesis to form two new A_s_ cells, or bona fide A_pr_ cells that will go on to form a chain of A_al_ cells. In rare cases (4%) EOMES was detected in some or all of the individual cells of GFRA1+ chains of A_al_ cells (Fig. 2F4, G and figure legend). In all instances, EOMES was co-expressed with PLZF (Fig. 2E, arrows), which marks all undifferentiated spermatogonia. EOMES was not detected in LIN28A-expressing cells (Fig. 2E), which we detect in a subset of A_s_ cells and in A_pr_ and A_al_ cells (*28*).

Because the EOMES+ population also expressed GFRA1, we hypothesized that these cells might be under GDNF influence (*23*). We therefore asked whether Sertoli-specific overexpression of GDNF affected this cell population. In whole-mount *Tg(Gdnf)* tubules we detected EOMES+ cells in clusters of PLZF+ and GFRA1+ cells (Fig. 2H, arrows). LIN28A stained only the peripheries of these cell clusters, and was specifically excluded from the EOMES+ cores (Fig. 2H, bottom row, asterisks). These data indicate that EOMES primarily marks a subpopulation of cells that are GFRA1+, PLZF+ and LIN28A-.

To determine if EOMES-expressing cells are totally absent in *lu/lu* testes, we immuno-stained *lu/lu* tubules for EOMES and GFRA1. Despite the significant decrease in *Eomes* RNA in whole testes, EOMES+ cells were detected in *lu/lu* mutants, (Fig. 2I and I’, arrows), suggesting that PLZF regulates the pool of EOMES+ cells, but does not directly regulate EOMES expression.

### EOMES+ cells contribute to steady-state spermatogenesis

One of the features of stem cells is their ability to maintain their population throughout adult life while giving rise to differentiating progeny. To study the stem cell behavior of EOMES expressing cells, we generated an inducible *iCreERt2/2A/tdTomato* knock-in allele (*Eomes*^*iCreERt2*^) at the *Eomes* locus to label and trace their progeny after tamoxifen treatment (Fig. 3A). Characterizing the mouse line for tdTOMATO expression, we observed single GFRA1+ tdTOMATO+ cells in whole mount seminiferous tubules from 4-week old males (Fig. 3B top row arrows). EOMES immunofluorescence marked the same cells as tdTOMATO (Fig. 3B’, top row), and as with EOMES itself, tdTOMATO was detected in a subpopulation of GFRA1+ cells (Fig. 3B’ bottom row, arrow). In testis sections, GFRA1+ tdTOMATO+ cells were detected within the seminiferous tubules (Fig. 3C - top) and along the basement membrane juxtaposed to peritubular myoid cells (Fig. 3C – bottom, arrowheads).

**Figure 3.**
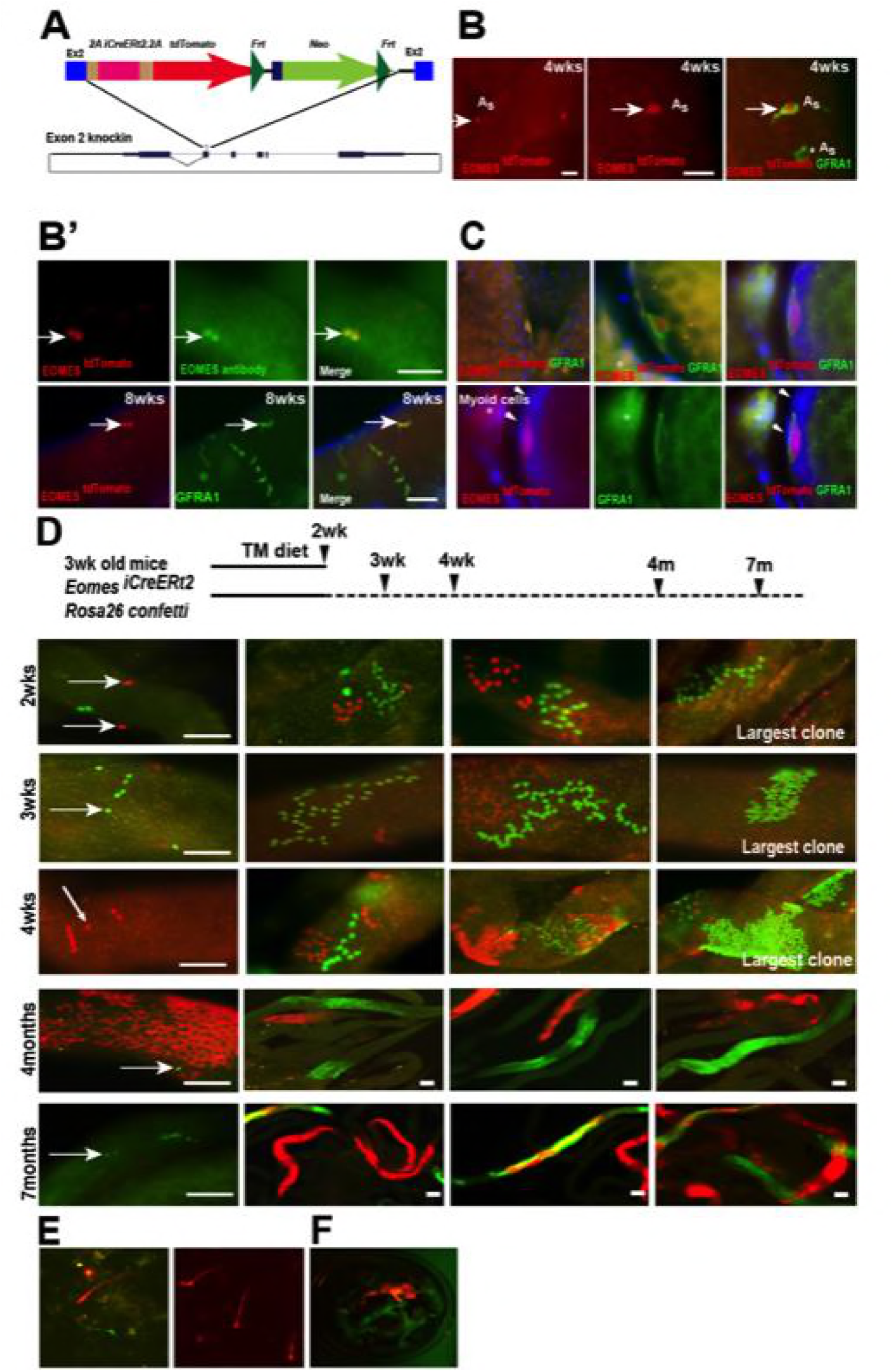
EOMES+ cells are long-term SSCs. (A) Schematic diagram of the *Eomes* locus on chromosome 9 and insertion cassette used to generate an *Eomes/2AiCreERT2/2A/tdTomato* knock-in allele. (B) GFRA1+ tdTomato+ A_s_ cell in testis of 4wk old mouse (arrow). GFRA1+ tdTOMATO-A_s_ cell (asterick). (B’) tdTOMATO is co-expressed with EOMES (top row, arrow). tdTOMATO is expressed in GFRA1+ A_s_ cells but not in chains of GFRA1+ A_al_ cells (bottom row).(C) tdTOMATO cells (Red), positive for GFRA1 (Green), are present within the seminiferous tubules. Top - An optical section of confocal image showing the localization of EOMES+ cell along the basement membrane. Bottom – Magnified view of GFRA1+ EOMES ^tdTOMATO+^ cells shown above. Arrowheads mark myoid cell (M) nucleus, which are negative for EOMES ^tdTOMATO^. * Background fluorescence in the interstitial space associated with GFRA1 staining. (D) Experimental flow chart showing the induction times and clonal analysis of *Eomes*^*iCreERt2*^ targeted cells labeled in mice crossed to Rosa26 Confetti reporter and fed a tamoxifen-enriched diet. Fluorescence images of whole-mount seminiferous tubules showing the different size of labeled clones. Representative clones at 2 (n=5, animals), 3 (n=4), and 4wks (n=3), and 4 (n=3) and 7 (n=3) months after tamoxifen induction. (E) Images of labeled sperm from the epididymis of mice 7months after tamoxifen induced recombination. (F) Fluorescence positive offspring born to WT female mated with ROSA26confetti: *Eomes-iCreERt2* 4months after tamoxifen-induced recombination at Confetti locus by *Eomes-iCreERt2*.

Next, we traced the progeny of *Eomes* ^iCreERt2^ cells during normal steady-state spermatogenesis after tamoxifen-induced recombination using the *R26R-confetti* reporter mice. Mice were fed a tamoxifen-enriched diet for 2 weeks and clone sizes were determined by the number of cells expressing a single fluorescent protein after recombination at the *R26R-confetti* locus and detected by fluorescence imaging of whole mount seminiferous tubules (Fig. 3D). One color clones indicate derivation from a single-labeled cell. We observed confetti-marked single cells and chains of 4, 8 and 16 by two weeks (Fig 3D, top row). The size of labeled clones increased with time (Fig. 3D, second and third row) and large labeled clones of differentiated germ cells were observed at 4 and 7 months after *iCre* activation (Fig. 3D). Importantly, we observed single-labeled cells at all time points analyzed (Fig. 3D, arrows), suggesting the single-labeled cells were either maintained for the entire duration or had undergone self-renewal and migrated away from its sister cells during the time the clones were traced. The labeled clones gave rise to labeled sperm detected in the epididymis of these mice (Fig. 3E), and fluorescence positive pups were born to *Eomes*-^*iCreERt2*^: *R26R-confetti* mice treated with tamoxifen when mated with WT C57BL6/J females (Fig. 3F). Labeled clones, or offspring, were not observed in the absence of TAM at any time points (Fig. S2). Together, these results demonstrate that EOMES-expressing cells contribute to long-term maintenance of A_single_ spermatogonia and to full-lineage differentiation to spermatozoa in mice.

### EOMES+ cells are resistant to busulfan treatment

To determine if EOMES+ cells are capable of regeneration after chemical insult, we specifically killed proliferating spermatogonia with busulfan and observed how EOMES+ and EOMES− cells survived the insult. We chose a low dose (10mg/kg body weight) that specifically kills proliferating A_pr_ and A_al_ spermatogonia (*35*). Seminiferous tubules from treated mice were immunostained for EOMES and PLZF every 3 days between 3 and 15 days after busulfan injection (Fig. 4A). During the entire 15-day time period the EOMES+ population remained unchanged (Fig. 4B). In contrast, by 9 days after treatment, the PLZF+ population had halved due to loss of A_pr_ and A_al_ cells. The population began to recover after 12 days (Fig. 4B). We also stained for EOMES and GFRA1. At day 0, ~20% of the total GFRA1-expressing cell population was EOMES+ (Fig. 4C). Comparison within the GFRA1+ population showed a selective increase of the EOMES+ fraction after busulfan treatment (Fig. 4C), suggesting that the EOMES− fraction was specifically lost.

**Figure 4.**
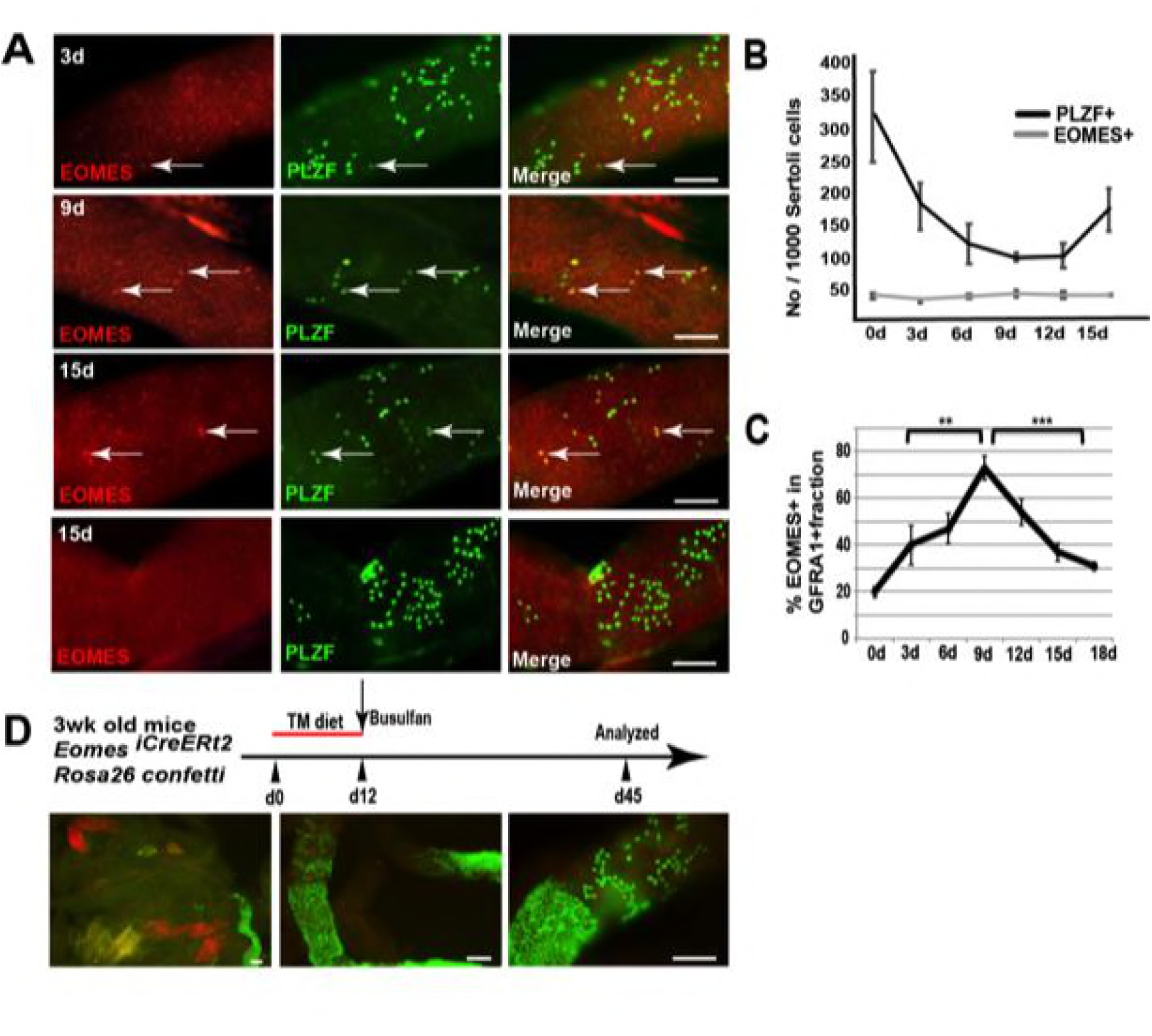
EOMES+ cells are resistant to busulfan and contribute in regeneration. (A) Whole-mount tubule immunostaining for EOMES and PLZF after low-dosage busulfan treatment (10mg/mL). (B) Cell counts from (A) show that EOMES+ cell numbers remain constant while PLZF+ cell numbers drop after busulfan treatment. (n = 3 per time point, 8 weeks old) (C) The EOMES+ fraction of GFRA1+ cells increased following busulfan treatment. **, p<0.005; ***, p<0.0001. (D) Schedule to assay the regeneration properties of EOMES+ cells. Three-week old *R26R-confetti*: *Eomes*^*iCreERt2*^ mice were put on a tamoxifen-enriched diet for 1 week, treated with busulfan, and analyzed on day 45. Images of whole mount seminiferous tubules (n = 5 mice) show labeled clones at different developmental stages during regeneration.

To assess if EOMES+ cells contribute to regeneration after chemical injury, 3wk old *Rosa26confetti: Eomes*-^*iCreERt2*^ mice were placed on a TM diet for 12 days to activate CreERt2, treated with a low dose of busulfan (10mg/kg body weight) on day 12, and then analyzed for labeled clones on day 45 (Fig. 4D). Whole mount seminiferous tubules showed labeled clones at varying stages of differentiation, supporting the conclusion that EOMES+ GFRA1+ PLZF+ spermatogonia are resistant to busulfan insult compared to their EOMES− counterparts, and that the EOMES+ population reconstitutes the lineage after chemical injury.

### EOMES+ A_s_ cells are slow-cycling spermatogonia

Proliferating cells are more susceptible to busulfan insult because the damaging alkylating activity crosslinks DNA during replication. We hypothesized that the resistance of the EOMES+ population to busulfan might be due to a difference in cell cycle rate compared to other spermatogonial populations. To test this, we used 5-ethynyl-2’-deoxyuridine (EdU) labeling to identify cells in S-phase, coupled with GFRA1 or EOMES immunostaining 24 hours after EdU injection (Fig. 5A). In WT tubules, only 10% of EOMES+ cells were labeled with EdU compared to 24% of the entire GFRA1+ population (Fig. 5B), suggesting that the EOMES+ subpopulation of GFRA1+ cells has a lower proliferation index.

**Figure 5.**
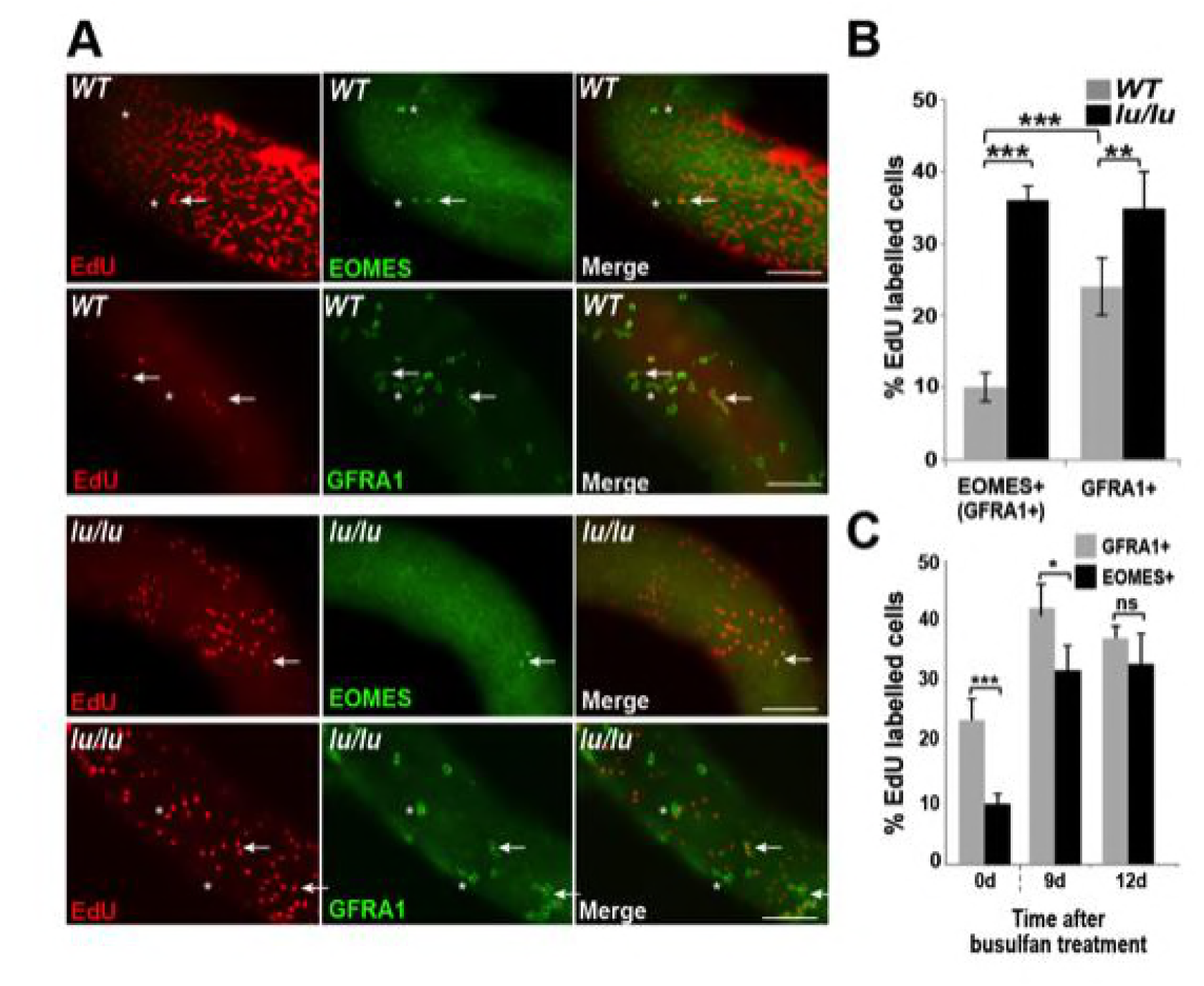
EOMES+ cells have a low proliferative index. (A) Whole-mount seminiferous tubules from 8-week old WT and *lu/lu* mice injected with EdU and immunostained 24 hours after injection. EdU+ EOMES+ or EdU+ GFRA1+ cells (arrows); EdU-EOMES+ or EdU-GFRA1+ cells (asterisks). (B) Quantification of the EdU labeling index in WT (n = 4 mice) and *lu/lu* (n = 5 mice) EOMES+ and GFRA1+ cells. GFRA1+ cells have a higher labeling index than EOMES+ cells in WT (p = 0.001). EOMES+ cells have a higher labeling index in *lu/lu* than in WT (p = 0.001). GFRA1+ cells have a higher labeling index in lu/lu than in WT (p = 0.005). (C) Increase in proliferation rate of EOMES+ and GFRA1+ cells during the recovery phase after busulfan treatment. EdU incorporation was assayed 24 hours post EdU injection (n = 3 mice per timepoint; *** p = 0.001; *p < 0.05).

In *luxoid* testes, we observed that EOMES+ cell numbers were greatly reduced. We speculated that progressive germ cell loss in the absence of *Plzf* could be due to an alteration in SSC cycling rates resulting in SSC exhaustion. To test this hypothesis, we injected EdU into *luxoid* mice and probed for cells in S-phase in conjunction with EOMES and GFRA1 immunostaining. Although fewer EOMES+ cells were present in *lu/lu* testes compared to WT, a greater fraction of the EOMES+ cells had incorporated EdU compared to WT (36% vs. 10%, Fig. 5A and B). Additionally, EdU labeling was significantly increased for the entire GFRA1+ population, but to a lesser degree (35% vs. 24%, Fig. 5B). These results indicate that EOMES+ cells have a higher than normal proliferative index in the absence of PLZF, leading to an overall increase in the number of total cycling GFRA1+ A_s_ spermatogonia.

To determine if the fraction of cycling EOMES+ cells changes during regeneration, mice were injected with EdU at 9 or 12 days post busulfan injection and assayed by immunofluorescence in whole mount seminiferous tubules. At both time points the EdU labeling index was increased in the EOMES+ and GFRA1+ populations (Fig. 5C) suggesting that busulfan is either directly inducing proliferation or the cells are responding to a decrease in SSC density.

### Single cell RNA sequencing of EOMES ^tdTOMATO+^ cells

The expression or function of a growing collection of genes have been linked to either SSC self-renewal or progenitor cell expansion (Table 1). To address if these SSC identity genes are expressed in EOMES+ cells, we performed single cell RNA sequencing (scRNAseq) on 897 fluorescently activated cell sorted (FACS) EOMES ^tdTOMATO+^ cells from 4wk old *Eomes-iCreERt2-tdTomato; Plzf ^+/+^* and 352 EOMES ^tdTOMATO+^ cells from *Eomes-iCreERt2-tdTomato; Plzf ^lu/lu^* mice (Fig. 6A). The mean number of expressed genes per cell in *Plzf*^*+/+*^ and *Plzf*^*lu/lu*^ was 3104 and 3230, while the mean number of transcripts with unique molecular identifiers was 7254 and 8134, respectively (Fig. S3).

**Table 1:** Genes reported to be expressed in SSCs or progenitor cells.

| Gene | Expression |  | Functional Assay | References |
| --- | --- | --- | --- | --- |
| ***Gfra1*** | A<sub>s</sub>, A<sub>pr</sub>, A<sub>al</sub> (4-8) | *Gfra1*-CreERT2 | Lineage tracing<br>Transplant | (7) |
| ***Bmi1*** | A<sub>s</sub> | *Bmi1*CreERT | Lineage tracing | (16) |
| ***Id4*** | A<sub>s</sub> | *Id4*-eGFP | Transplant | (15) |
| ***Plzf*** | A<sub>s</sub>, A<sub>pr</sub>, A<sub>al</sub> (4-16) | Antibody staining | Transplant | (26) |
| ***Lin28a*** | A<sub>s</sub>, A<sub>pr</sub>, and A<sub>al</sub> (4-16) | Antibody staining | Transplant | (28) |
| ***T*** | A<sub>s</sub> | *T*(nEGFP-CreERT2) | Lineage tracing | (17) |
| ***Eomes*** | A<sub>s</sub> | *Eomes*-iCre ERT2<br>tdTomato | Lineage tracing | This manuscript |
| ***Nanos2*** | A<sub>s</sub>, A<sub>pr</sub> | Antibody staining | Knockout | (11) |
| ***Nanos3*** | A<sub>s</sub>, A<sub>pr</sub>, A<sub>al</sub> | Antibody staining | Knockout | (8) |
| ***Neurog3*** | A<sub>al</sub> (4-16) | *Ngn3* CreERT | Lineage tracing | (6) |
| ***Pax7*** | A<sub>s</sub> | *Pax7*-CreERT2 | Lineage tracing | (14) |

**Figure 6.**
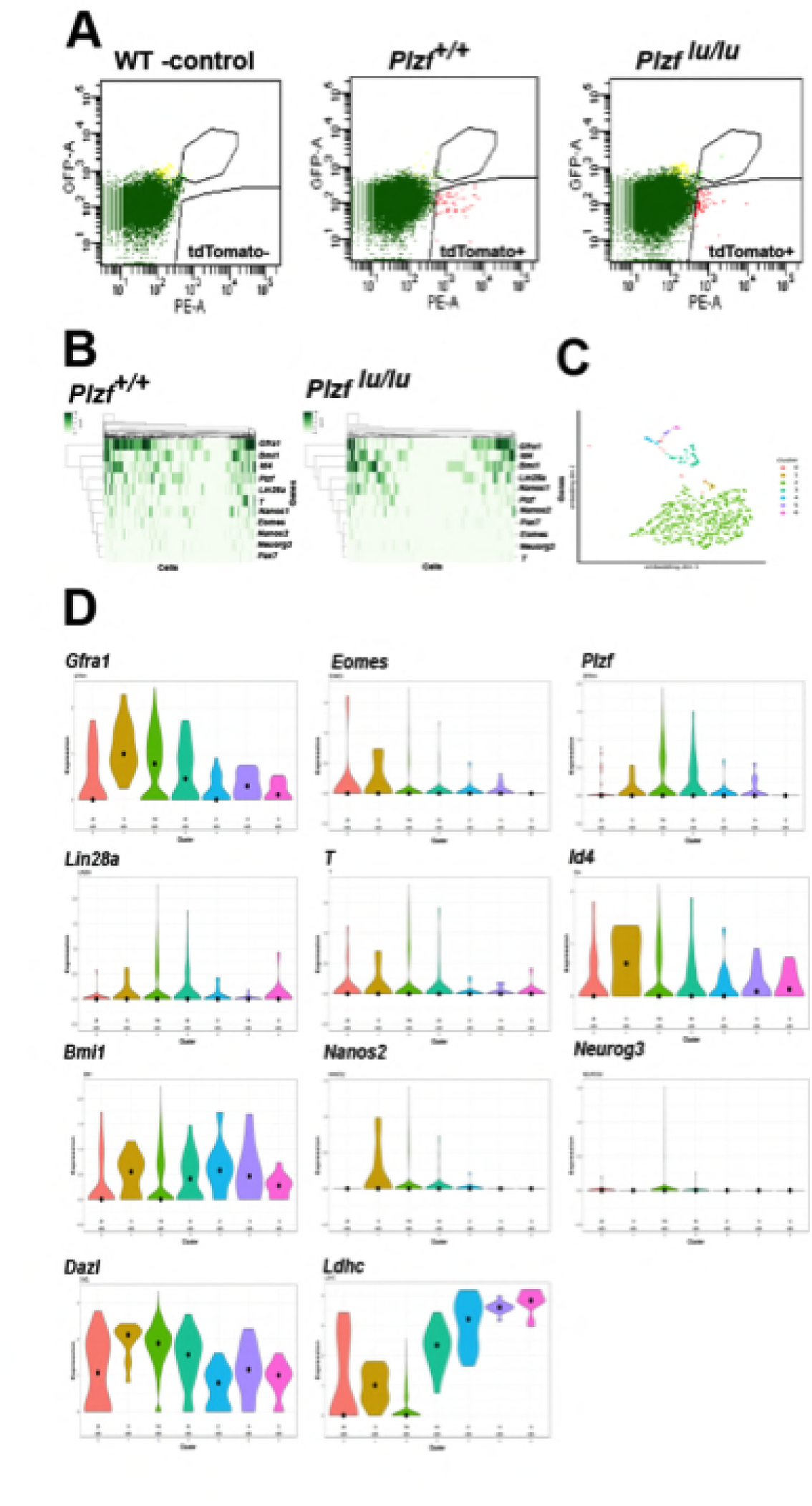
Single cell RNA sequencing of EOMES ^tdTOMATO^ cells. (A) FACscan profile of EOMES ^tdTOMATO+^ cells sorted from *Plzf*^*+/+*^ and Plzf ^*lu/lu*^ mutant testis. The boxed region in the upper right indicated cells with background autofluorescene that were excluded from the analysis. (B) Heat map and dendogram of SSC identity and progenitor genes expressed in EOMES ^tdTOMATO^ cells from *Plzf*^*+/+*^ and Plzf ^*lu/lu*^ mutant testis. (C) Clusters generated from WT cells. Cellular expression profiles of 110 genes were embedded into a 2dimensional latent space using UMAP (see supplemental information for details). Cells in cluster 0 could not be confidently placed in any of the other 6 clusters. (D) Violin plots showing the mean expression of each SSC identity gene per cluster in cells sorted from *Plzf*^*+/+*^ testes.

An expression heat-map of the 11 identity genes indicates that the majority of the cells express *Gfra1*, *Bmi1*, *Id4* and *Plzf*, while fewer cells express *T*, *Eomes*, *Nanos*, *Neuorg3* and *Pax7 (*Fig. 6B). The most striking finding is the considerable degree of heterogeneity in expression across the entire scRNAseq data set. None of the cells expressed all of the genes, while 90% (807/897) of cells expressed at least one of the genes and 73% (655/897) expressed two or more of the genes. *Gfra1* was detected in 566/897 cells (63%), while PAX7 was detected in only 10/897 cells (1.11%). *Neurog3* (*36*), which is expressed in early progenitor cells, was detected at relatively low levels in 27/897 cells (3.01%), suggesting that *Neurog3* is expressed in some SSCs, or that the sorted cells include early progenitor cells. *Eomes* transcripts were detected in 85/897 (9.48%) EOMES ^tdTOMATO+^ cells. The low number of cells could be due to the inherent low expression of *Eomes* relative to other expressed genes, early progenitor cells that are tdTOMATO positive but are no longer expressing *Eomes* transcript, or the cell cycle stage in which the cells were collected. Together, the heterogeneity in transcript detection in the scRNAseq data mirrors previously reported heterogeneity in protein expression as detected by immunofluorescence (*8*).

To determine the relationship between the cells expressing the identify genes, cluster analysis was performed on the 100 most differentially expressed genes, plus 10/11 identity genes, from *Plzf*^*+/+*^ sorted cells*. Pax7* was excluded from the analysis because its normalized mean expression was too low to be meaningful. The bulk of the cells clustered together (cluster 2) as a group of 740 cells (Fig. 6C). The remaining cells were defined by a separate group of 5 different smaller clusters. Cells assigned to cluster 0 could not be confidently assigned to any cluster. The expression of *Dazl,* a marker of germ cells (*37*), at moderate to high levels in all clusters, confirms that each cluster represents germ cells (Fig. 6C). The expression of *Ldhc*, which is highly restricted to germ cells and whose protein is first detected in early pachytene spermatocytes (*38*), is expressed at highest levels in clusters 3-5, which are represented by a few cells, suggesting that *Ldhc* mRNA is expressed at low levels in SSCs or early progenitor cells. Although *Gfra1* was detected in all clusters, its highest mean expression was in cluster 1, as was *Id4* and *Bmi* (Fig. 6D). Although cluster 1 contained only 13 cells, *Eomes*, *T* and *Nanos2* were expressed in a higher proportion of cells (e.g. 23%-38% in cluster 1 compared to 9%-15% in cluster 2) than in any other cluster. Nonetheless, the presence of multiple clusters expressing different identity genes suggests that EOMES ^tdTOMATO+^ sorted cells represent a collection of cells with different transcriptional profiles and supports the conclusion that SSCs are transcriptionally heterogeneous.

To determine the impact of loss of *Plzf* on the expression of the identity genes, we computed the mean expression level of each gene across all cells in *Plzf ^+/+^* and *Plzf*^*lu/lu*^ mice. In *Plzf*^*lu/lu*^ mice, the mean expression of the identity genes *Bmi1*, *Id4, Pax7, Nanos2* and *Neurog3* were unchanged compared to *Plzf +/+* mice, while the mean expression of *Gfra1*, *Eomes* and T were significantly decreased (Fig. 6B and Table 2). In *Plzf*^*lu/lu*^ mutants, the fraction of cells expressing *Gfra1* was reduced from 63.10% in *Plzf*^*+/+*^ cells to 38.07% in *Plzf*^*lu/lu*^ mutants. *Eomes* and *T* were only expressed in (1.14%) of *Plzf*^*lu/lu*^ mutant cells, compared to 9.48% and 14.94%, in *Plzf*^*+/+*^ cells, respectively, suggesting that mutation of *Plzf* differentially affects a subpopulation of SSCs expressing *Gfra1*, *T* and *Eomes*.

**Table 2:**
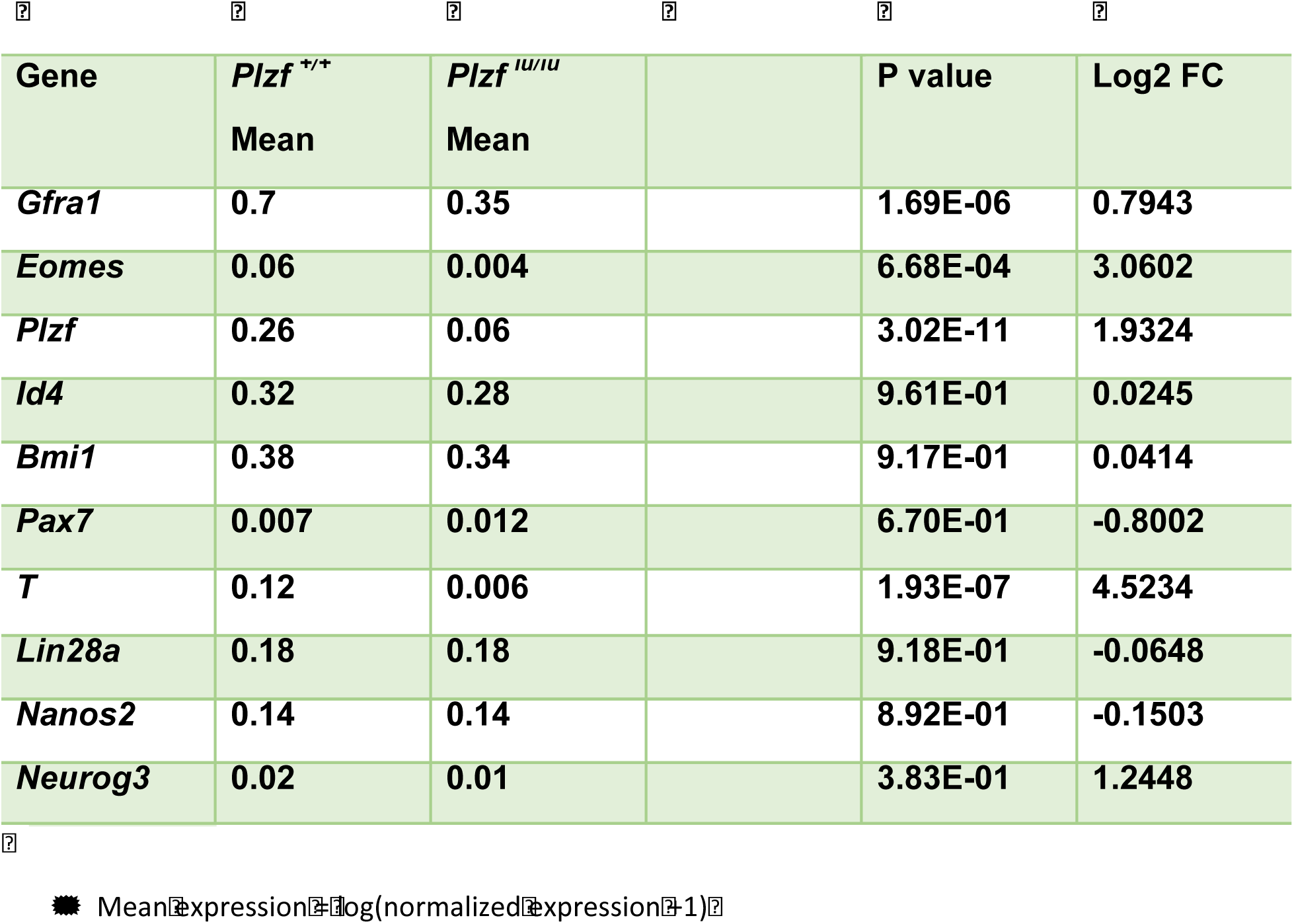
Mean expression* of SSC identity transcripts in *Plzf*^*+/+*^ and *Plzf*^*lu/lu*^ mutant cells.

Given that ID4-EGFP has previously been reported to be restricted to a subpopulation of A_s_ cells (*39*), that it has been proposed to mark the ultimate pool of SSCs (*39*), and that we did not detect significant changes in *Id4* transcript in EOMES ^tdTOMATO+^ sorted cells in *Plzf ^lu/lu^* mutants, we investigated the expression of ID4 by immunofluorescence. Using a rabbit polyclonal antibody, which detects a protein of the correct molecular weight by western blotting in extracts prepared from WT testes but not in extracts from *Id4^-/-^* testes (Fig. 7A), we detected ID4 in PLZF+ A_s_, A_pr_, and A_al_ cells (Fig. 7C). Because this staining pattern differed from that previously reported for an *Id4-EGFP* transgene, we performed double-staining for ID4 and LIN28A. We again found that we could detect ID4 in chains of A_al_ cells that were also positive for LIN28A (Fig. 7D). ID4 immunostaining was not detected in seminiferous tubules of two different *Id4 ^-/-^* mutants (Fig. 7E). We assayed for co-expression of tdTOMATO and ID4 and found that 40% of the tdTOMATO+ A_s_ cells were also positive for ID4 (Fig 7F, upper row). We again detected ID4 in chains of A_al_ cells (Fig. 7F, bottom row), none of which were positive for tdTOMATO. In aggregate, we determined that only 20% of ID4+ cells were positive for tdTOMATO. To determine if *Id4* was required for expression of EOMES, we performed immunofluorescence on testis from *Id4^-/-^* mutants. EOMES was detected in GFRA1+ cells (Fig. 7G), consistent with the detection of *Eomes* RNA by rtPCR in *Id4^-/-^* mutants (Fig. 7B). We conclude that ID4 is expressed in a subpopulation of EOMES+ cells, but the majority (80%) of the ID4+ population is EOMES-.

**FIGURE 7.**
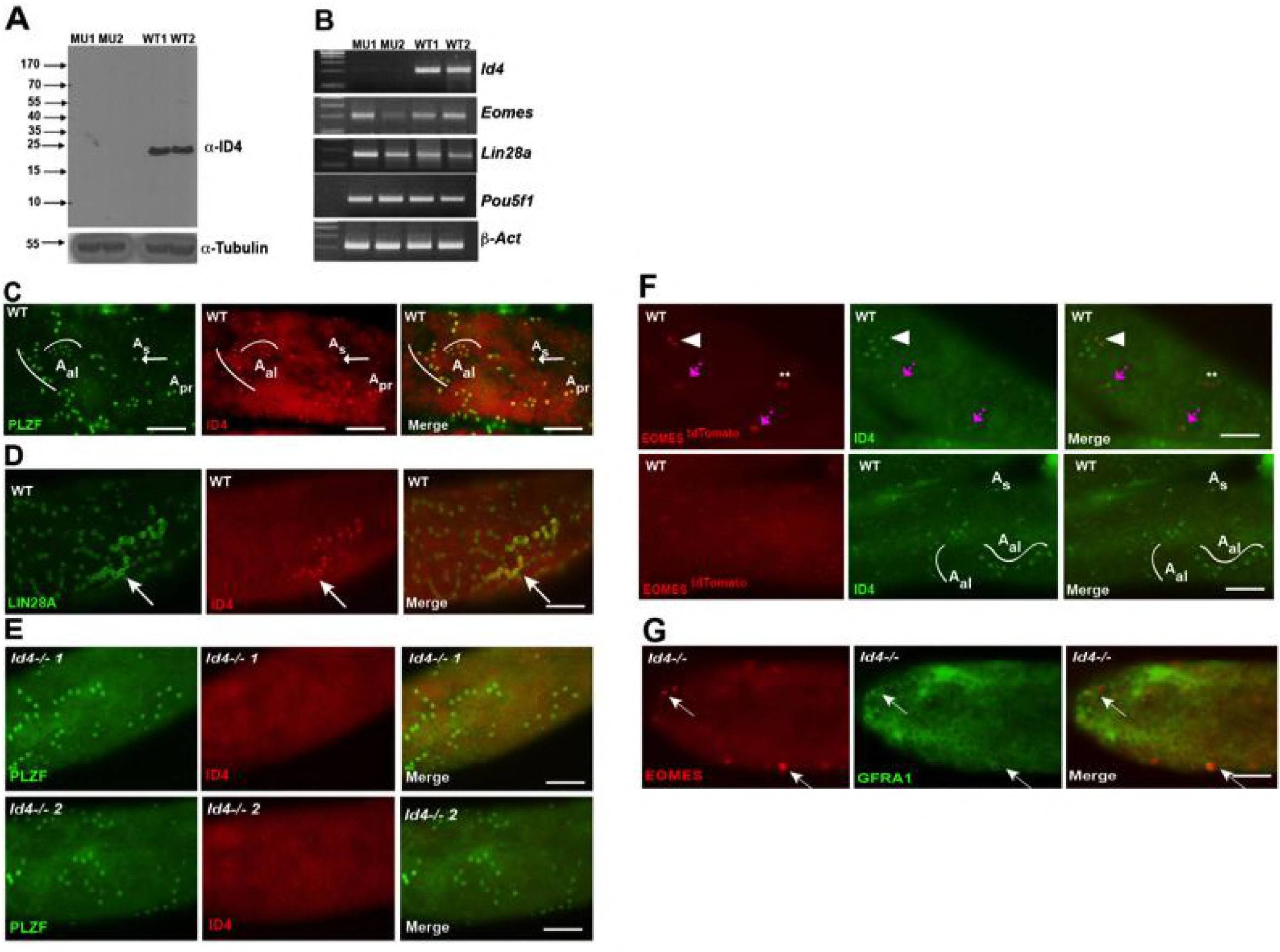
ID4 is expressed in A_s_, A_pr_ and A_al_ spermatogonia. A)Western blot of total testis protein lysate from 2 WT and 2 *Id4^-/-^* mutant mice probed with ID4 antibody. For loading control, western blot was probed with α-tubulin antibody. (B) RT-PCR using total RNA from two *Id4^-/-^* mutant and 2 WT mice for *Eomes, Lin28a, Pou5f1 and β-actin*. (C) Detection of PLZF and ID4 by immunofluorescence in whole mount seminiferous tubules. Both ID4 in PLZF+ were detected in A_s (arrow)_, A_pr_, and A_al_ spermatogonia in WT adult mouse testis. (D) ID4 protein was co-localized in chains of LIN28A+ A_al_ spermatogonia (arrow). (E) ID4+ cells are not detected in whole mount tubules from prepared from *Id4^-/-^* mutant mice. (F) Top row, examples of tdTOMATO/ID4 double-positive cells (arrowhead), tdTOMATO+ ID4-(**), and two adjacent cells where one is tdTOMATO+ and the other is tdTOMATO/ID4 double-positive (magenta arrow). Bottom row, examples of tdTOMATO-ID4+ A_s_ and A_al_ chains of 4 and 8 cells. (G) EOMES+ GFRA1+ cells are present in *Id4*^-/-^ mutants (arrow).

## DISCUSSION

Exploiting a transgenic overexpression model for GDNF in *Plzf ^+/+^* and *Plzf lu/lu* mice, we discovered a rare population of EOMES-expressing spermatogonia. Lineage tracing studies during steady state spermatogenesis and during regeneration demonstrated that EOMES+ cells are SSCs. EOMES+ cells cycle more slowly than the bulk of GFRA1+ cells, and in *Plzf ^lu/lu^* mice, both populations cycle more frequently. The significant loss of *Eomes-* and *T-* expressing cells between *Plzf ^+/+^* and *Plzf ^lu/lu^* suggest that EOMES and T represent a subpopulation of GFRA1+ cells that are uniquely depleted in *Plzf ^lu/lu^* mice.

The concept of a slow-cycling or long-term SSC in rodents, referred to as the A_0_ “reserve stem cell”, was first proposed 50 years ago (*40*). In retrospect, the authors were most likely distinguishing between rapidly proliferating A1-A4 spermatogonia and undifferentiated A_s_-A_al_ spermatogonia. The A_0_ cell was never discovered, and the field turned towards the ‘A_s_’ model, which stipulated that A_s_ cells were the functional SSC population that both self-renewed and proliferated irreversibly into differentiated cell subtypes (*2, 41*). Early 3[H]-thymidine incorporation studies hinted at the existence of both rapid-cycling and long-cycling A_s_ SSCs (*42*). However, similar studies in Chinese hamsters did not support the earlier studies in rats (*43*).

With our discovery of slow-cycling EOMES+ cells under PLZF-mediated proliferative control, we now suggest that the A_s_ population is heterogeneous in its cycling status. In our working model, there are two subtypes of SSCs under steady-state conditions: a slow-cycling (A_s-SC_) long-term stem cell that expresses EOMES and a more rapid-cycling (A_s-RC_) short-term stem cell that is EOMES negative. Both populations express GFRA1 and PLZF. This functional hierarchy ensures long-term tissue homeostasis of the stem cell compartment and continued spermatogenesis. We propose that during steady state conditions, the A_s-RC_ cells divide regularly to both self-renew and differentiate. Although these cells populate the spermatogonia lineage and support the normal cycle of the epithelium, we suggest their capacity is limited. Upon depletion or reduction of A_s-RC_ cells in a given region of the epithelium, A_s-SC_ cells are recruited and convert to EOMES-A_s-RS_ cells. Key to this model is the limitation of cycling in A_s-SC_ cells by PLZF, which prevents stem cell exhaustion and the progressive germ cell loss observed in *luxoid* mutants. By extension, our data suggest that the failure of *luxoid* mutant germ cells to successfully colonize after transplantation is a consequence of their severely depleted population of EOMES+ A_s-SC_ cells and the altered proliferative index of the entire population of GFRA1+ cells. This paradigm could extend to the regulation of germline stem cells in humans, as PLZF is expressed in human SSCs (*44*), which include a proposed quiescent A_s_ subpopulation.

Overexpression of GDNF led to clusters of tightly packed nests of undifferentiated spermatogonia. Cells in the center of the nests tended to be EOMES+ LIN28A-while cells at the periphery were EOMES-LIN28A+. We speculate that the significance of the geometry of the nests could be a consequence of increased paracrine signaling, perhaps driven by intercellular coupling of GDNF with its co-receptor GFRA1/RET, or the physiological state in the center of the nests, that favors SSC self-renewal.

In addition to expression in A_s_ cells, EOMES was occasionally detected in one of two adjacent GFRA1+ cells. The clones, which we counted as A_pr_, could have also been two lineage-independent, but adjacent, A_s_ cells. Alternatively, the two cells could be daughter cells, suggesting that EOMES expression can be asymmetrically inherited. Interestingly, in rare cases EOMES was expressed heterogeneously in chains of GFRA1+ A_al_ cells, again suggesting that the fate of cells with clones of A_al_ is not uniform and supporting the possibility that cells within chains are subject to reversion to SSCs following fragmentation (*6*).

### Relation of EOMES cells to other SSC markers

Single-cell RNA sequencing of EOMES ^tdTOMATO+^ cells from *Plzf ^lu/lu^* mutants suggests that EOMES-expressing A_s_ cells constitute a unique population of A_s_ cells. In recent studies, other GFRA1+ A_s_ subpopulations have also been identified. *Pax7* lineage-marked cells are capable of repopulating germ cell-free testes in transplants and are resistant to busulfan insult. (*14*). However, unlike EOMES+ cells, PAX7 cells are rapid cycling, and *Pax7* transcript was detected in only 10/897 EOMES ^tdTOMATO+^ cells, suggesting the possibility that they correspond to rapid-cycling EOMES- GFRA1+ A_s_ SSCs. A *Bmi1 ^CreERT^* allele has also been shown to mark SSCs (*16*). We detected *Bmi1* transcript in EOMES ^tdTOMATO+^ cells, and long-term labeling of SSCs has been observed in *Bmi1* lineage tracing experiments, however, the mean expression of *Bmi1* was similar between *Plzf ^+/+^* and *Plzf ^lu/lu^* cells, suggesting that *Bmi1* expression is not restricted to EOMES+ cells.

An *Id4-gfp* allele also marks a subset of A_s_ spermatogonia (*12, 13, 15, 45*) and recent studies have shown that transplantable SSC activity resides in the ID4-GFP-High population (*39*). Because ~40% of EOMES+ cells are ID4+, and *Id4* transcripts were co-expressed in 49 of the 85 *Eomes+* cells, it is likely that there is substantial overlap in the population of cells marked by EOMES, ID4-GFP-High, and T, which has recently been shown to also be expressed in a subpopulation of A_s_ cells with SSC activity (*17*). However, our data suggest that the *Id4-gfp* transgene may not reflect the full expression pattern of the ID4 protein itself. Although we detected ID4 in about 40% of the EOMES+ cells, the majority of the ID4+ cells (80%) were EOMES- and included A_pr_ and A_al_ cells. Differences in half-lives of the ID4 and EGFP proteins, or differences in translational control elements in their respective mRNAs, may be responsible for the discordance in their expression.

At face value our results are inconsistent with the fragmentation model for SSC self-renewal (*6, 7*). However, it is possible that the *Ngn3* and *GFRa1* lineage tracing studies preferentially labeled the equivalent of our EOMES-GFRA1+ rapid-cycling SSCs, which we propose function as short-term SSCs. It also remains a possibility that EOMES-A_s_, A_pr_ and A_al_ cells can de-differentiate to EOMES+ A_s_ slow-cycling SSCs following fragmentation.

### Function of PLZF

To date, it is unclear how PLZF mechanistically regulates SSC self-renewal, as it is expressed in the entire undifferentiated spermatogonia population. Several studies have provided *in vitro* evidence for a role of PLZF in cell cycle regulation. PLZF associates with CDC2 kinase activity *in vitro* (*46*) and when overexpressed in myeloid progenitors, PLZF decreases proliferation by reducing entry into S-phase (*47*). Furthermore, PLZF suppresses growth of 32Dcl3 cells *in vitro* by repressing transcription of the cyclin A2 gene and other cyclin-dependent complexes involved in the G_1_/S transition (*48, 49*). Our studies show that PLZF is critical for maintaining SSCs in a low proliferative state and suggest that it regulates the cycling status of SSCs. Analogous to the hematopoietic system, which contains both slow-cycling long-term (LT)-HSCs, and rapid-cycling short-term HSCs, (*50–52*), and where over-cycling of LT-HSCs can lead to stem cell exhaustion and consequent loss of differentiated cell types (*53*), we propose that loss of PLZF results in proliferative exhaustion of SSCs.

## Acknowledgements

We are extremely grateful to Dr. Matthew Havrda for providing us with *Id4*^-/-^ mice, to Dr. Dayana Krawchuk for her help in manuscript preparation and to The Single Cell Biology Lab at The Jackson Lab for Genomic Medicine.

## Author contributions

M.S. designed, executed and wrote the manuscript. A.S. analyzed the RNAseq data. H.E.F and D.B. contributed to experimental design and execution. W.F.F. performed the scRNAseq analysis. R.E.B contributed to experimental design, interpretation of the data and writing of the manuscript.

## Funding

This work was supported by a grant from NICHD/NIH (HD042454UW, to R.E.B.) and from NCI (CA34196, to The Jackson Laboratory) in support of The Jackson Laboratory’s shared services.

## Supplementary Information

**Figure S1.**
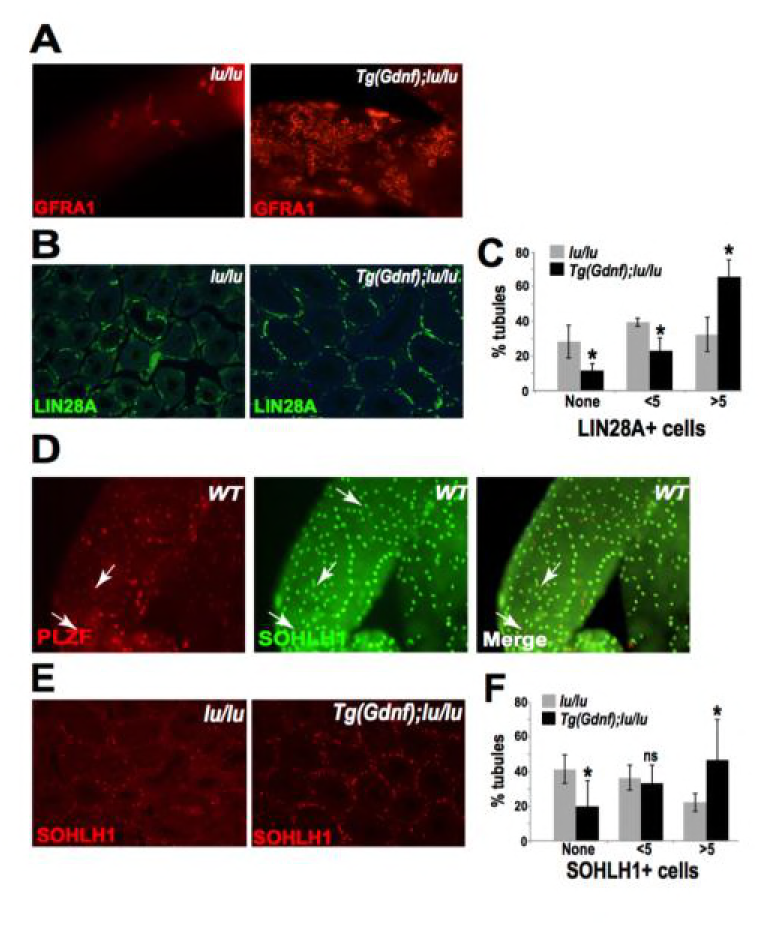
(Related to Fig. 1) Sertoli-cell overexpression of GDNF increases the SSC population in *Plzf-mutant luxoid mice*. (A) Immunostained whole-mount tubules from 12-week old *Tg(Gdnf);lu/lu* mice show large clusters of GFRA1+ cells compared to *lu*/*lu*. (B) Representative LIN28A immunostaining of *lu/lu* and *Tg(Gdnf);lu/lu* tubule paraffin sections. (C) Quantification of immunostained sections as in (B) show 70% more tubules with 5 or more LIN28A+ cells in *Tg(Gdnf);lu/lu* compared to *lu*/*lu* (n=4). *, p<0.05. (D) In WT SOHOLH1 expression is detected in A_al_ cells transitioning to A1 and differentiating cells A2-A4, absent from A_s_, A_pr_, and A_al_ cells. (E) Representative SOHLH1 immunostaining of *lu/lu* and *Tg(Gdnf);lu/lu* tubule paraffin sections. (F) Quantification of immunostained sections as in (E) show a significant increase in the number of tubules with 5 or more SOHLH1+ cells in *Tg(Gdnf);lu/lu* compared to *lu*/*lu*. (n=4). *, p<0.05, ns, not significant.

**Figure S2.**
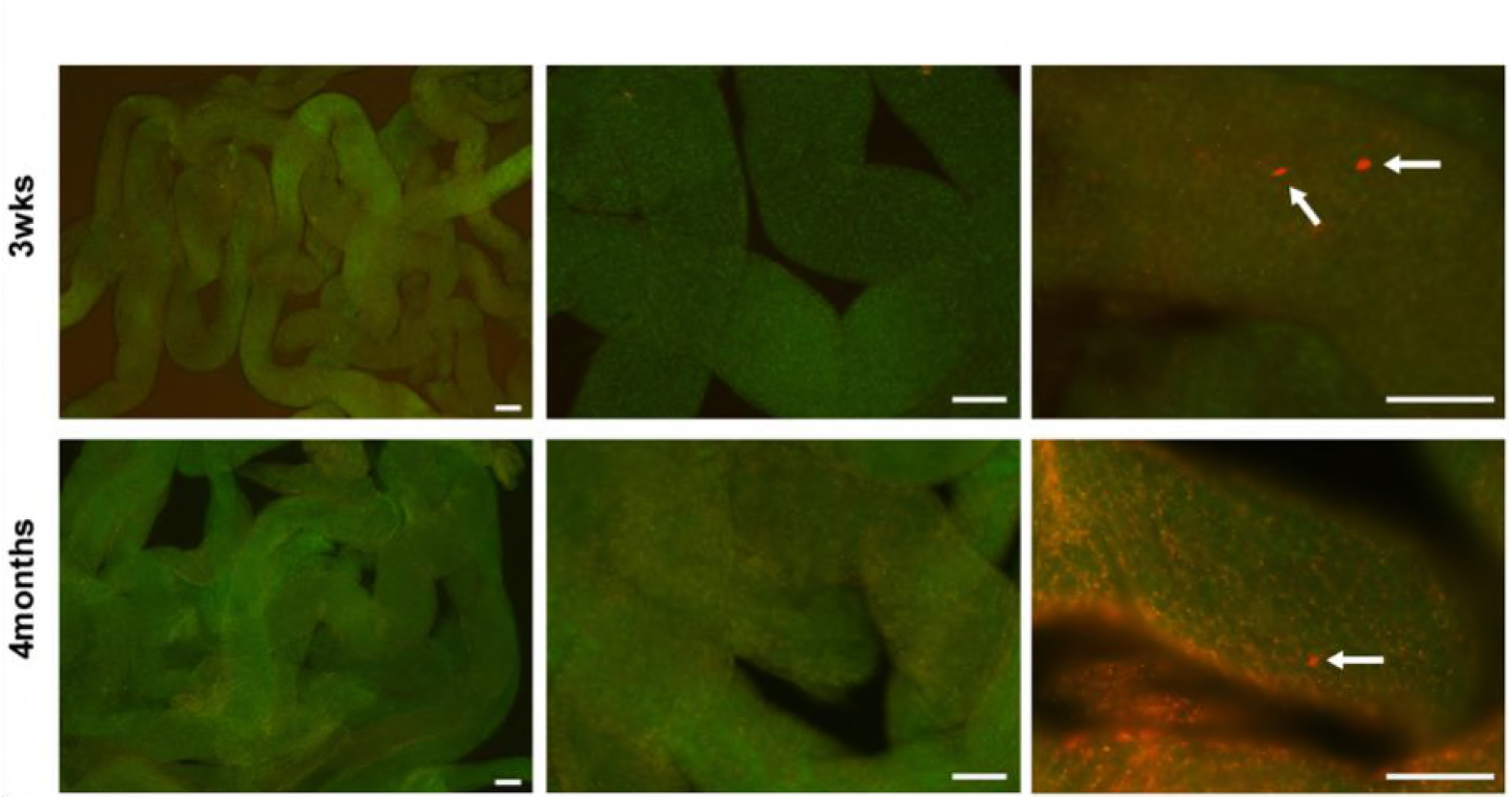
(Related to Fig. 3.) No TAM controls for lineage tracing of *Eomes iCreERt2* targeted cells in mice crossed to Rosa26 Confetti reporter on a normal (no tamoxifen) diet. Representative images at 3wks and 4mos post completion of TM diet fed to experimental cohorts shown in Fig. 3. Arrows indicate tdTOMATO+ cells. Scale bar = 100μm

**Figure S3.**
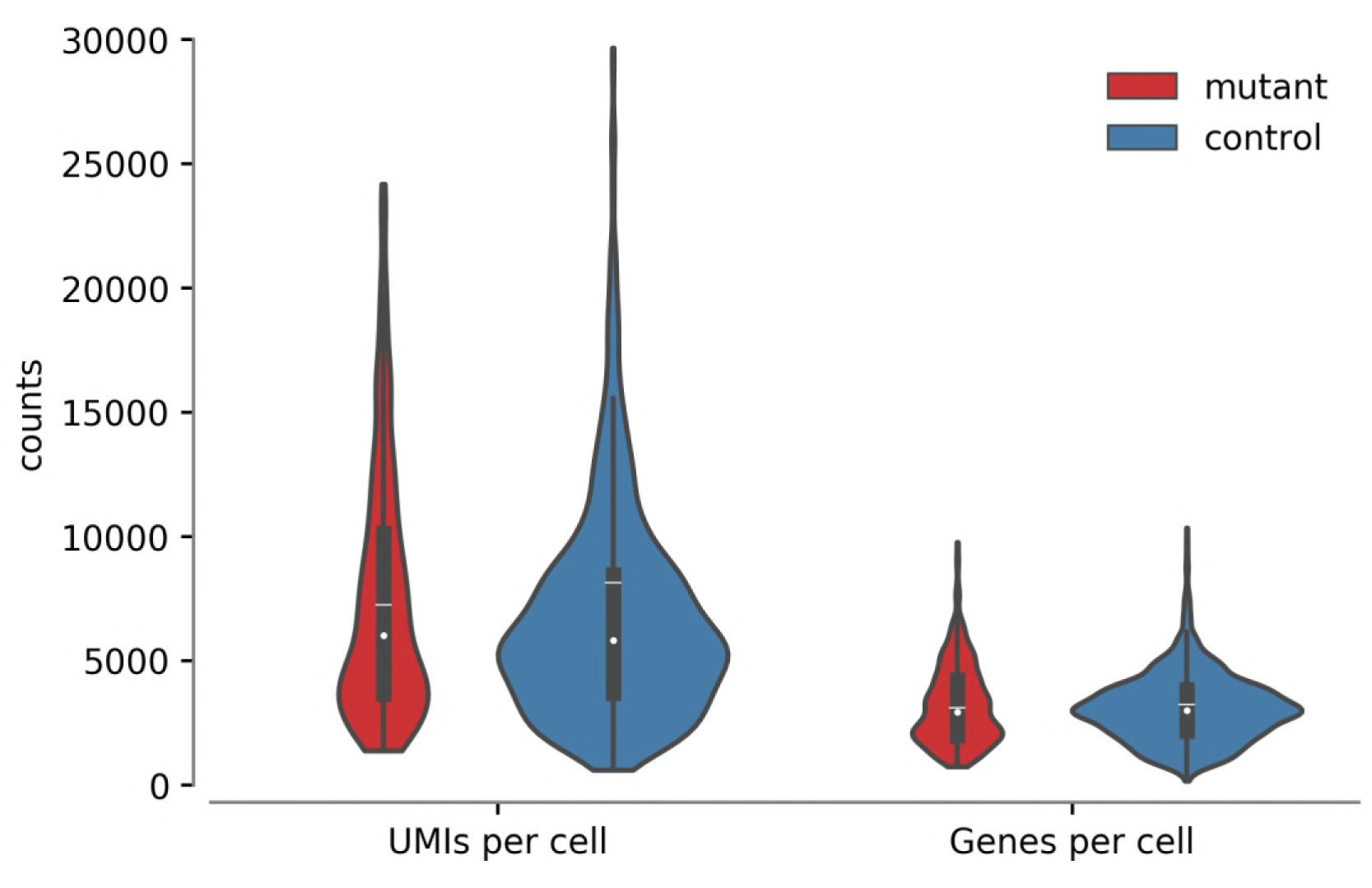
(Related to Fig. 6) Distribution of unique molecular identifiers (UMIs) and genes per cell. Median (white dot), mean (white line), and interquartile range (dark box) are shown for each of the four distributions, with control (*Plzf ^+/+^; Eomes-iCreERt-tdTomato+*) in blue and mutant (*Plzf ^lu/lu^; Eomes-iCreERt-tdTomato+*) in red.

## MATERIALS AND METHODS

### Animals

For *Plzf ^lu/lu^*, we used the Green’s luxoid or ‘luxoid’ (as referred to here) *Zbtb16^lu^*/J allele from The Jackson Laboratory. B6.Cg-*Gt(ROSA)26Sor*^*tm6(CAG-ZsGreen1)Hze*^/J (CAG(ZsGreen) mice (*54*) were obtained from the The Jackson Laboratory (JR # 007906). *Id4*^-/-^ mice (*55*) were provided by Mathew Havrda (Dartmouth University). All animals were housed in a barrier facility at The Jackson Laboratory. All experimental procedures were approved by the Jackson Laboratory Institution Animal Care and Use Committee (ACUC) and were in accordance with accepted institutional and government policies.

### Generation of Tamoxifen inducible *iCre ERt2-tdTomato* knock-in into the *Eomes* locus

The BAC clone RP23-235G22 was used to clone the *2A:CreERt2-FRT-PGK-Neor-FRT* cassette into exon 2 of mouse *Eomes* gene. Linearized vector was electroporated into B6N-JM8 ES cells from B6N mouse strain. Chimera were generated using B6(Cg)-^Tyrc-2J^/J the albino mice (are C57BL/6J mice carrying a mutation in the tyrosinase gene). The chimeras were crossed to B6N parental strain to generated F1 progeny. The reporter mice used to assay for Cre recombination were *B6.129P2-Gt(ROSA) 26 ^Sortm1(CAG-Brainbow2.1)Cle^/J* also known as R26R-confetti conditional mice stock number # 017492 from Jackson laboratory. For activation of *iCre* the mice were fed TD.130859 (TAM diet). The food was formulated for 400mg tamoxifen citrate per kg diet that would provide ~40mg tamoxifen per kg body weight per day.

### Testes weight and histology

Mice were genotyped with specific primer sets and assessed for body weight and testes weight at each time point analyzed. For histological analysis, testes were fixed overnight in Bouin’s and paraffin-embedded sections were stained with PAS after dewaxing.

### Antibodies

Antibodies were purchased from the following companies. Mouse anti-PLZF, antibody (Santa Cruz Biotech Inc., Santa Cruz, CA); Rat anti-GFRA1 (R&D Systems, Minneapolis, MN); Rat anti-LIN28A (gift from Dr. Eric G. Moss, Dept. of Mol. Biology, UMDNJ, New Jersey); Rabbit anti-SOHLH1 and anti-EOMES from (Abcam, Cambridge, MA), and Rabbit monoclonal anti-mouse /human ID4 antibody (# BCH-9/#82-12) from BioCheck, Inc. Foster City, CA.

### Whole mount immunostaining

Seminiferous tubules were dispersed in PBS by removing the tunica from the testes. They were rinsed with PBS and fixed in 4% paraformaldehyde (Electron Microscopy Sciences, Fort Washington, PA) for 4-6h at 4°C. For PLZF, LIN28A, and SOHLH1 antibodies, the tubules were permeabilized with 0.25% NP-40 in PBS-T (0.05% Tween in PBS) for 25 min at RT before blocking in 5% normal goat serum. The tubules were then incubated with primary antibodies and incubated overnight (GFRA1 and EOMES) at RT or 4°C for PLZF, LIN28, and SOHLH1. The next day the tubules were washed three times with PBS-Tween20 for 5 min, and incubated with secondary antibodies conjugated to Alexa Fluors (Molecular Probes) for 1 h at room temperature. After washing in PBS-Tween20, the tubules were mounted in VectaShield® with DAPI (Vector Laboratories, Burlingame, CA) and imaged using a Nikon Eclipse E600 epifluorescence microscope equipped with an EXi Aqua Camera from Q-Imaging (Surrey, BC, Canada).

### Proliferation assay

To assay the proliferative index of spermatogonial stem cells, mice were injected intra-peritoneally with 50mg/kg body weight of EdU. EdU staining was performed using the Click-iT^Tm^ EdU imaging Kit (Invitrogen, Carlsbad, CA) according to manufacturer’s protocol. Briefly, seminiferous tubules were dissected 24 hrs after EdU injection and first probed with GFRA1 or EOMES antibodies before staining for EdU. Tubules were incubated with Click-iT ^TM^ reaction cocktail containing buffer, CuSO_4_, Alex Fluor 594 Azide, and reaction additive buffer for 30 min at room temperature. Tubules were then washed in PBS-Tween20 buffer for more than 2 hours and then mounted in Vectashield mounting media (Vector Laboratories Inc., Burlingame, CA) with DAPI and imaged on a Nikon Eclipse E600 epifluorescence microscope equipped with an EXi Aqua Camera from Q-Imaging (Surrey, BC, Canada).

### Quantitative Real Time (qRT)-PCR

Total RNA was prepared from testes using the PureLink^TM^ Kit (Ambion). qRT-PCR was performed using SYBR^®^ Green Master Mix with gene-specific primer sets in a One-step RT-PCR reaction (Applied Biosystems) on an equal amount of total RNA from testes of each genotype and analyzed using ABI 7500 system sequence detection software (Applied Biosystems, Carlsbad, California). Arcturus^TM^ PicoPure^TM^ RNA kit from Applied Biosystems (Waltham, MA) was used to isolated RNA from the FACsorted cells.

### mRNA sequencing and analysis

An equal amount (4µg) of total RNA from testes of 4 month-old mice was used for sequencing using the TruSeq RNA Sample Prep Kit v2. The libraries were amplified by 15 PCR cycles and quantitated using a Kapa Biosystems Quantification Kit. The quality of each library was assessed on a Bioanalyzer Agilent DNA 1000 Chip. The libraries were then normalized to 2nM and then pooled into 1 sample. The pooled samples were loaded onto 3 lanes of a V3 Flow Cell and then sequenced on the Illumina HiSeq 2000 (100bp paired-end run). RNAseq data for all samples were subjected to quality-control check by NGSQCtoolkit v 2 (*56*), and samples with base qualities ≥ 30 over 70 nucleotides (100 bp reads) were used in the analysis. Orphaned reads after the quality-control step were used as single-ended reads in the downstream analysis. Read mapping was carried out using *TopHat v 2.0.7* (*57*) with supplied annotations at parameters (–read-mismatches 2, –read-gap-length 3 and –read-edit-dist 3) against the mouse genome (build-mm9). Bamtools v 1.0.2 (*58*) was used to calculate the mapping statistics. *HTSeq-count* script was used at default parameters (http://www-huber.embl.de/users/anders/HTSeq/doc/count.html) to count the reads mapped to known genes for both orphaned and paired-end reads, and final counts were merged prior to differential expression analysis. Expression differences between *Tg(Gdnf)* and *Tg(Gdnf);lu/lu* mice were determined by using *edgeR v 2.6.10* (*59*) (*exactTest* function was used to determine the differences in expression). Genes with FDR ≤ 0.05 and Absolute log_2_ fold change ≥ 1 were considered differentially expressed.

### Regeneration of SSCs after busulfan treatment

A single dose of 10mg/kg body weight of busulfan was injected intraperitoneally in WT C57BL/6 adult mice. Three mice were analyzed at each time point. Their testes were processed for whole-mount immunofluorescence. For a percentage of GFRA1 population, 300-500 cells were counts for each mouse. The EOMES+ and PLZF+ cells were counted over equal length of seminferous tubules, for each mouse. The data are presented as the number of cells ± SE per 1,000 Sertoli cells.

### Flow Cytometry and single cell sequencing

Single cell suspensions from *Plzf ^+/+^* and *Plzf ^lu/lu^* testes carrying the *Eomes-tdTomato* knockin allele were prepared using a two-step collagenase and trypsin enzymatic digestion procedure. Cells were filtered first with an 40um nylon membrane followed by a 25um filter. DNase I was added to the cell suspension to prevent clumping. Cells were washed once in PBS and finally suspended at a concentration of 1-2×10^6^ cells/ml, analyzed and sorted on Becton Dickinson FACSVantage cell sorter. Cells from littermate mice that were negative for *Eomes-^tdTomato^* were used to gate the positive staining cell population and DAPI was used to gate out the dead cells. Single cells were flow sorted into individual wells of a Bio-Rad hard shell 384 well plate. The plate was immediately transferred and stored in a −80C freezer. Custom designed Drop-Seq barcodes (see below) from Integrated DNA Technologies (IDT) were delivered into each well of each 384 well plate. All primers in one well shared the same unique cell barcode and billions of different unique molecular identifiers (UMIs). An Echo 525 liquid handler was used to dispense 1ul of primers and reaction reagents into each well in the plate for the cell lysis and cDNA synthesis. Following cDNA synthesis, the contents of each well were collected and pooled into one tube using a Caliper SciClone Liquid Handler. After treatment with exonuclease I to remove unextended primers, the cDNA was PCR amplified for 13 cycles. The cDNA was fragmented and amplified for sequencing with the Nextera XT DNA sample prep kit (Illumina) using custom primers (see below) that enabled the specific amplification of only the 3′ ends. The libraries were purified, quantified, and sequenced on an Illumina NextSeq 500.

Barcode oligo primer: 5’-AAGCAGTGGTATCAACGCAGAGTACJJJJJJJJJJJJNNNNNNNN TTTTTTTTTTTTTTTTTTTTTTTTTTTTTTVN-3’

Custom primer sequence: 5’-AATGATACGGCGACCACCGAGATCTACACGCCTGTCCGCGGAAGCAGTGG TATCAACGCAGAGT*A*C-3’

Custom read 1 sequence: 5’-CGGAAGCAGTGGTATCAACGCAGAGTAC-3’

### Single cell sequencing analysis

Cells derived from the mutant and control samples were clustered separately. Genes with fewer than 2 counts in fewer than 2 cells were removed from the digital expression matrix and the expression was normalized by relative library size (computed as total counts per cell divided by median counts per cell) and log transformed. 100 genes with the largest variance / mean ratio and 10 genes used for cell type verification (*Gfra1*, *Eomes*, *Id4*, *T*, *Lin28a*, *Plzf*, *Bmi1*, *Pax7*, *Nanos2* and *Nanos3*) were used to subset the expression matrix for dimensionality reduction and clustering. Cellular expression profiles at these 110 genes were embedded into a 2-dimensional latent space using UMAP (McInnes L and Healy J. Uniform manifold approximation and projection for dimension reduction. ArXiv e-prints 2018, version 0.2.1), and clusters of cells were identified using hierarchical density-based spatial clustering (HDBSCAN, Campello Ricardo JGB et al., Hierarchical density estimates for data clustering, visualization, and outlier detection. ACM transactions on Knowledge Discovery from Data (TKDD) 2015; McInnes L, Healy J, and Astels S. Hierarchical density based clustering. Journal of open Sourse Software (JOSS) 2017, version 0.8.12). Cells assigned to cluster 0 are those which could not be confidently assigned to any cluster.

